# Communication pathway analysis within protein-nucleic acid complexes

**DOI:** 10.1101/2025.02.14.638259

**Authors:** Sneha Bheemireddy, Roy González-Alemán, Emmanuelle Bignon, Yasaman Karami

## Abstract

Inter-residue communication forms a vast and intricate network that underpins essential biological processes such as catalysis, gene expression, and cell signaling. Allostery, a crucial phenomenon where distant regions of a macromolecule are energetically coupled to elicit functional responses, operates through these intricate communication networks within macromolecular complexes. Despite the pivotal role of nucleic acids in these networks, their contributions to allostery remain largely overlooked. To address this gap, we developed ComPASS, a large-scale computational method designed to study communication networks in protein-protein and protein-nucleic acid complexes. Recognizing the significance of dynamics in the communication of macromolecules, our approach leverages molecular dynamics (MD) simulation data to extract inter-residue key properties, including dynamical correlations, interactions, and distances. These properties are integrated to construct a weighted communication network that comprehensively represents dependencies among amino acids and nucleotides. Using ComPASS, we uncovered distinct mechanisms of signal transmission in diverse macromolecular systems. In Cysteinyl-tRNA synthetase, the central domain was found to mediate the coordination between substrate recognition and enzymatic activity, ensuring functional precision. In the LacI repressor, allosteric communication occurs through interface pathways within the dimer, effectively linking ligand sensing to DNA binding. For the Type IIF restriction endonuclease Bse634I, structural communication across dimer and tetramer interfaces was crucial for specific DNA recognition. In the liver X receptor, a key helical region was identified as a bridge connecting ligand-binding events to DNA interactions. Finally, our analysis with ComPASS aligned with previous literature, confirmed the role of H2A L1 loops in mediating communication across histone interfaces and coordinating interactions between structural domains in nucleosome complexes. ComPASS is available as an open-source tool, maintained at https://github.com/yasamankarami/compass. By offering an integrated framework for studying communication networks, ComPASS advances our understanding of conformational dynamics, particularly within protein-nucleic acid complexes.

## Introduction

The study of macromolecular functions is shifting from focusing on isolated protein structures to constructing integrative models that encompass the complexity of biological assemblies. This paradigm shift contributes to the growing recognition that static representations of individual proteins, while valuable, cannot capture the dynamic interplay and conformational changes that drive biological processes. Recent advancements, such as AlphaFold3 [1], have revolutionized static structure prediction, achieving unprecedented accuracy for diverse proteins. However, AlphaFold3 and similar approaches are inherently limited to static models and cannot account for the dynamic behavior and communication essential for understanding the functions in large macromolecular complexes. Molecular dynamics (MD) simulations have emerged as a powerful tool to address this gap, providing atomistic insights into the temporal and spatial dynamics of biomolecules. MD simulations facilitate the exploration of protein flexibility and conformational changes that underlie biological functions. These simulations generate data with potentials to reveal critical insights into macromolecular dynamics. However, effectively extracting and integrating this information into coherent models remains a crucial challenge, showing the need for novel approaches to fully harness the dynamic data provided by MD simulations.

Communication networks in macromolecular complexes are fundamental to the coordination and regulation of biological processes, enabling the transmission of signals and information across different regions of the complex. Among these, allosteric pathways represent a critical subset, where perturbations at one site are transmitted through the structure to modulate the activity of a distant functional site [2]. The study of allosteric pathways aids in understanding processes such as signal transduction [3], allosteric regulation [4], and coordinated conformational changes [5], revealing the basis of response by macromolecules to environmental stimuli and functional specificity. This is also instrumental in drug design [6], enabling the development of therapeutics that target dysfunctional parts of the network. Advancements in computational and experimental techniques have attempted for the detailed mapping of these networks, uncovering the complex interplay of structural and dynamic elements that facilitate allosteric regulation.

Over the past decade, the understanding of allostery and communication networks has evolved significantly. Various computational tools have been developed to analyze molecular communication networks, often adopting a site-specific perspective to elucidate local interactions. Existing methods leverage either static structural data or dynamic conformational information obtained from MD simulations or normal mode analysis. Methods utilizing dynamic data frequently focus on dynamic correlations among residues to investigate allosteric mechanisms. These include interaction-based tools such as ProteinLens [7], PyInteraph2.0 [8], and NRIMD [9]; distance-based methods like DyNetAn [10] and SPMWeb [11]; correlation and covariance-based tools including MDiGest [12] and PSNTools [13]; and entropy and information theory-based approaches exemplified by Allopath [14]. While these tools provide valuable insights, majority of them are constrained to a single property or interaction type, limiting their ability to deliver a holistic perspective on allostery. While effective for targeted studies, such approaches may often overlook the broader context of communication pathways within entire complex. Comprehensive techniques that integrate data across the entire macromolecular complex are essential for capturing the intricacies of intra-complex communication. To address these limitations, integrative methodologies have emerged, combining multiple dynamic and structural features to study the communication network comprehensively. One such approach, COMMA [15], integrates five residue-level dynamic properties—local dynamical correlations, minimum distances, communication propensities, non-covalent interaction strengths, and secondary structure characteristics—to analyze conformational ensembles and derive communication pathways. Here, we propose **ComPASS**, a method designed to study communication networks in macromolecular complexes from a dynamic perspective. ComPASS extends the scope of previous methods by accommodating both protein-protein and protein-nucleic acid (protein-NA) complexes, offering a versatile and comprehensive tool to investigate communication pathways and functional relationships in diverse biological systems.

ComPASS is designed to extract and analyze key dynamic properties from MD simulations, offering a comprehensive framework for studying macromolecular communication networks. These properties include generalized correlations between residues, non-covalent interactions across the complex, communication propensities, and inter-residue distances derived from MD trajectories. By integrating MD data analysis with network-based approaches, ComPASS constructs intricate communication networks that enable the identification of communication pathways within macromolecular complexes. This approach addresses limitations of traditional methods that rely on static structures or use only one property, and are constrained to protein-protein interactions by broadening the scope to include protein-NA complexes. By capturing the temporal and spatial dimensions of macromolecular communication, ComPASS provides deeper insights into allosteric regulation and other functional mechanisms. Its ability to process large-scale MD datasets efficiently overcomes computational bottlenecks, enabling researchers to investigate complex biological systems more effectively. To demonstrate its capabilities, we applied ComPASS to various complexes derived from the Allosteric database (ASD) [16], highlighting its utility in unraveling intricate communication networks and functional mechanisms in diverse biomolecular contexts. ComPASS is freely available at https://github.com/yasamankarami/compass.

## Materials and Methods

### Dataset preparation

We selected five systems of interest, comprising four protein-NA complexes from the ASD database and the nucleosome core particle. For the complexes chosen from the ASD database we used the following filters: containing nucleic acid residues, residues annotated as allosteric, resolved structure with a corresponding PDB ID, and residues identified as being involved in allostery. This criteria yielded four complexes: *i)* E. coli cysteinyl-tRNA synthetase (CysRS) bound to tRNACys, *ii)* the Lac Repressor (LacI) bound to ONPG in its repressed state, *iii)* the Type IIF restriction endonuclease Bse634I (Bse634I) complexed with its cognate DNA, and *iv)* the heterodimer of retinoid X receptor *α*-liver X receptor *β* (RXR*α*-LXR*β*) bound to DNA. We considered the biological assemblies and removed the ligands for further analysis. For the last systems, there were two missing loops (K213-T223, Y249-E251) in RXR*α* and one missing loop (Y152-P164) in LXR*β*, which were modelled using Modeller [17]. We also analyzed three nucleosome complexes that have been previously investigated to explore communication pathways mediated by histone H2A variants [18]. They include: *i)* a canonical nucleosome (1KX5), *ii)* a variant of this complex with a mutation in the L1 loop sequences, replacing ^38^NYAE^41^ with ^38^HPKY^41^ as found in the macroH2A variant (1KX5_*L*1_), and *iii)* a nucleosome complex with the H2A.Z variant (1F66). A fourth nucleosome system featuring the 601 Widom sequence (NCP601) was also included in the study to take advantage of extensive MD data available from a previous study [19] and to further scrutinize possible DNA sequence effects on the DNA-histones communication pathways. **Table 1** summarizes the details of the studied systems.

**Table 1:**
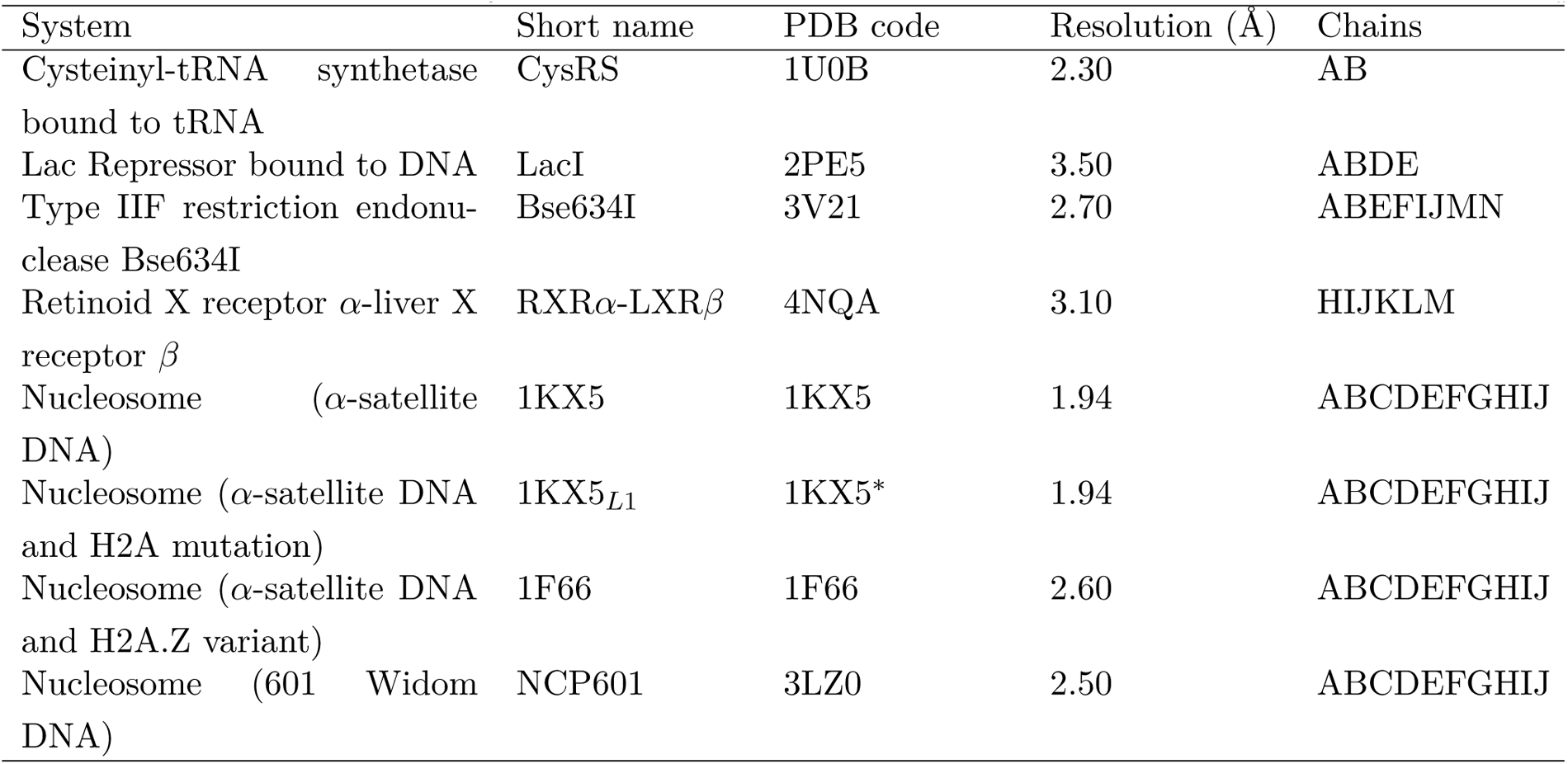
The list of studied systems. ^*∗*^ With the ^38^NYAE^41^ to ^38^HPKY^41^ mutation on the L1 loop.

### Setup of MD simulations

The following MD protocol was used for CysRS, LacI, Bse634I and RXR*α*-LXR*β*. The environment of the histidine was checked using MolProbity [20]. The systems were then prepared using the CHARMM-GUI interface [21–23] and simulations were carried out with GROMACS 2023.4 [24]. We used the Amber ff14SB force field [25] for proteins, the OL3 force field [26] for RNA, the BSC1 force field [27] for DNA, and the TIP3P water model [28]. Each system was solvated in a dodecahedron box with a minimum distance of 12 Å between the solute and the box edges, and counterions (Na^+^, Cl^*−*^) were added to reproduce physiological salt concentration of 150 mM. First, we performed 20000 steps of minimization using the steepest descent method keeping only protein backbone atoms fixed to allow protein side chains to relax. Then, the system was equilibrated for 125 ps at constant volume. For every system, three replicates of 500 ns, with different initial velocities were performed in the NPT ensemble. The temperature was kept at 310 K and pressure at 1 bar using the Parrinello-Rahman barostat with an integration time step of 2.0 fs. The Particle Mesh Ewald method [29] was employed to treat long-range electrostatics, and the coordinates of the system were written every 100 ps. MD for the three nucleosome systems 1KX5 (*α*-satellite sequence NCP from PDB 1KX5), 1KX5_*L*1_ (*α*-satellite sequence NCP from PDB 1KX5 with mutated H2A L1 loop) and 1F66 (*α*-satellite sequence NCP with the H2A.Z variant from PDB 1F66) were set up and performed as explained by Bowerman and Wereszczynski [18] in order to allow for a direct comparison between the pathways predicted by ComPASS and their results. The MD ensembles of the 601 Widom NCP taken from a previous study [19] consisted in three replicates amounting for a total of *∼*15 *μ*s. The simulation details are reported in **Supplementary Table S1**.

#### Stability of the trajectories

Standard analyses of the MD trajectories were performed using the *gmx* module of GROMACS 2023.4. The root mean square deviation (RMSD) of the backbone atoms (*Cα, C, N, O*) from the initial frame were recorded along each replicate (**Supplementary Fig. S1**). The by-residue root mean square fluctuations (RMSF) were calculated on the backbone atoms (*Cα, C, N, O*), with respect to the first frame (**Supplementary Fig. S2**).

### Network parameters

#### Generalized correlations

The generalized correlation matrix is calculated from MD trajectory data to capture non-linear dependencies between residues. We start by extracting the positions of the *n* alpha carbon (CA) or C5′ atoms across all frames of the trajectory. For each pair of residues *i* and *j*, we compute the covariance matrix and extract the mutual information matrix, which is then transformed into a generalized correlation coefficient to better capture the complex interactions inherent in macromolecular complexes. This transformation involves using mutual information as an intermediary, ensuring that the resulting matrix reflects both linear and non-linear correlations. The generalized correlation coefficient *GCC*_*ij*_ between the *i*-th and *j*-th atoms is calculated as:

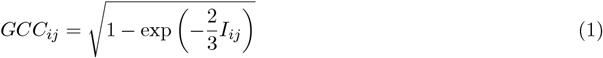

where *I*_*ij*_ is the mutual information.

#### Interaction propensity

ComPASS identifies hydrogen bonds, salt bridges, and close contacts between all pairs of residues for each frame of the MD trajectory. For each type of interaction, it produces a matrix that displays the frequency of those interactions along with an additional matrix that captures the occurrence of any detected interactions. Tests were carried out to ensure that the geometries of the hydrogen bonds aligned with the output of the Baker-Hubbard algorithm [30] implemented in MDTraj [31]. Additionally, close contacts were validated to closely correspond with those detected using HBPlus [32]. Similarly, the salt bridges were successfully compared with the results of the corresponding VMD plugin [33].

#### Communication propensity and distance matrices

The communication propensity (CP) matrix is derived from the variance of the distances between the pairs of residues on the MD trajectory. For each pair of residues *i* and *j*, we compute the *CP*_*ij*_ following our previous work [15]:

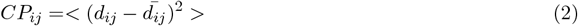

where *d*_*ij*_ is the distance between the *Cα* atoms of residues *i* and *j* and 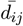 is the mean value computed over all the conformations. For each pair of residues *i* and *j*, we calculate the minimum distance between all atoms in each frame and then average these distances across all frames. This distance matrix provides a measure of the spatial proximity of residues throughout the simulation.

#### The adjacency matrix

The adjacency matrix, a critical component for network representation, was constructed by integrating the *generalized correlation matrix*, the *interaction frequency matrix*, and the *communication propensity matrix*. These matrices were first flattened into a combined matrix of dimensions *n* × 3*n*, where *n* represents the number of residues. Principal Component Analysis (PCA) was then applied to reduce the dimensionality of the combined matrix, retaining the top two principal components. This transformation resulted in a reduced matrix of dimensions *n ×* 2. Subsequently, the pairwise distances between the rows of this PCA-transformed matrix were calculated to generate a pairwise distance matrix of dimensions *n × n*. From this, a covariance matrix was computed, which captures the relationships between residue pairs. The decomposition of the eigenvalues was performed on the covariance matrix, and the two principal components (eigenvectors) corresponding to the largest eigenvalues were selected. These steps culminated in three *n × n* matrices that encapsulate key residue-residue relationships, forming the basis for the final adjacency matrix used in network analysis.

#### Network Construction

The nodes of the network are derived from MD trajectories, where the *n* alpha carbon or C5’ atoms of the given macromolecular complex are considered. Edges are constructed by applying a distance cutoff to the average minimum distance matrix. Although the user has the option of selecting a custom cut-off point for edge construction, the default value is set to 5Å. Finally, the adjacency matrix, derived from the interaction and correlation data, is used as a weight to construct a weighted graph that reflects the strength of interactions between residues.

### Analysis of network parameters

The constructed network was analyzed using several common network parameters to assess its topology and communication properties. All graph-theoretical descriptors were computed using Networkx. Detailed instructions on the execution and interpretation of the ComPASS Python code and the results are reported in **Supplementary methods**.

#### Computation of Communities

The constructed network was analyzed to identify functionally correlated residues, which were then classified into distinct communities. We used the Leiden algorithm for community detection due to its efficiency in handling large networks or graphs [34]. The Leiden algorithm optimizes the modularity of the network and is particularly effective for large-scale systems, making it ideal for macromolecular complexes. The resulting communities correspond to highly interconnected functional modules within the protein-NA network, highlighting groups of residues that may act in concert during various functional processes.

#### Identification of Cliques

Cliques represent fully connected subgraphs, where each node is directly connected to every other node in the subgraph. In the context of our network, cliques highlight tightly regulated, cohesive groups of residues that have high mutual interactions. We used networkx pymol package to identify subgraphs within the 10Å interaction graph. This approach allowed us to capture groups of residues that are likely to play crucial roles in maintaining the structural integrity and functional dynamics of the macromolecular complex.

#### Computation of Shortest Paths

Shortest paths represent the most efficient routes for communication between residues in the network. To compute these paths, we used Dijkstra’s algorithm, which is ideal for finding the shortest paths in weighted graphs. We calculated the shortest paths between every pair of residues in the network and then ranked them based on the weight of the pathways. The top 10% of the shortest paths, identified by their weight, were considered the most significant pathways in the given network. These paths provide valuable insights into how residues communicate and how information is transferred within the complex.

#### Computation of Optimal Paths

To further explore the communication pathways between specific residues, we computed the optimal paths between a given source and target residue using Yen’s algorithm[35]. This algorithm is particularly useful for finding multiple shortest paths between two nodes, which can reveal alternative communication routes. Given the source and target residues, as well as a user-defined number of requested paths, Yen’s algorithm computes the optimal paths that best reflect the most efficient or alternative routes for information transfer. These optimal paths provide deeper insights into the communication dynamics of the macromolecular complex.

#### Identification of Key Residues

To pinpoint crucial residues within the network, we considered several centrality measures: betweenness centrality, closeness centrality, and degree centrality. These centrality measures help to identify residues that act as key mediators in the network, either by controlling the flow of communication (betweenness), being well-connected to other residues (degree), or being important in terms of their proximity to other residues (closeness). Based on these measures, we identified the top 5% of residues as crucial players in the network, which are likely to be involved in important regulatory functions or allosteric interactions.

## Results

To investigate communication networks and allosteric mechanisms in macromolecular complexes, we developed ComPASS, a computational framework that integrates multiple dynamic properties extracted from MD simulations. As shown in **Fig. 1** and detailed in the **Methods section**, ComPASS constructs a weighted communication network based on dynamical correlations, non-covalent interactions, and distances. By systematically analyzing this network, we identify key structural and functional features, including communication pathways, communities, cliques, and allosteric hotspots. To illustrate the broad applicability of ComPASS, we applied it to five distinct macromolecular systems, each representing different modes of allosteric regulation. These case studies highlight how diverse biomolecular systems leverage communication networks to regulate their function. In the following sections, we present detailed analyses of each system, demonstrating how ComPASS captures key allosteric features and provides mechanistic insights into macromolecular communication.

**Figure 1:**
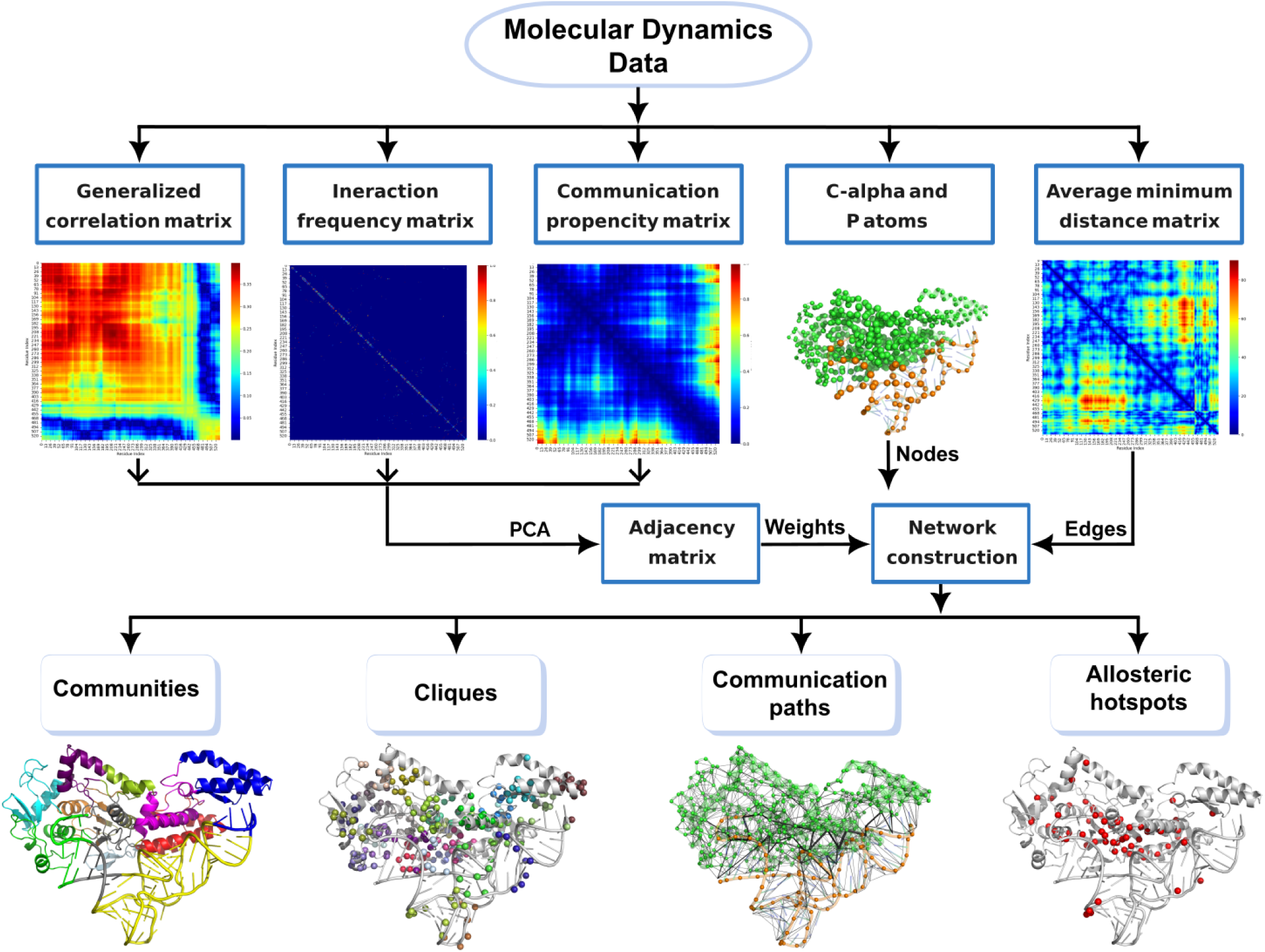
The workflow of ComPASS. The sequential steps are shown, starting from the extraction of properties from MD trajectory data to the construction and analysis of the final communication network.

### Core domain bridges amino acid recognition and catalysis in Cysteinyl-tRNA Synthetase

Aminoacyl-tRNA synthetases catalyze the attachment of amino acids to the 3’ end of their cognate tRNAs, ensuring accurate aminoacyl-tRNA synthesis for genetic decoding. In these enzymes, the recognition of the tRNA anticodon triplet is allosterically linked to catalytic efficiency at the active site, despite being physically separated by *∼*70 Å [36]. This communication involves dynamic, indirect, and direct readout mechanisms that cannot be fully captured by static crystal structures. To elucidate the functional dynamics of CysRS, we applied ComPASS to the generated MD trajectories, providing a detailed map of the networks underpinning aminoacylation specificity. As shown in **Supplementary Fig. S3A**, the communication network within the complex predominantly involves C*α*–nucleotide connections, underscoring the critical role of protein-RNA interactions in facilitating allosteric communication within this system. **Fig. 2A** illustrates the top pathways identified across the complex, with pathway weights used to rank their importance. This suggests that RNA binding plays a critical role in mediating communication within the protein, facilitating the transfer of information between the anticodon recognition domain and the active site. Specifically, the top pathways link the amino acid recognition site to the active site through the protein-RNA interface at the core domain. The results are in agreement with the literature, where extensive interaction between the helix bundle domain and the anticodon binding domain was shown to contribute to structural rigidity in the presence of tRNA, facilitating allosteric communication [36].

**Figure 2:**
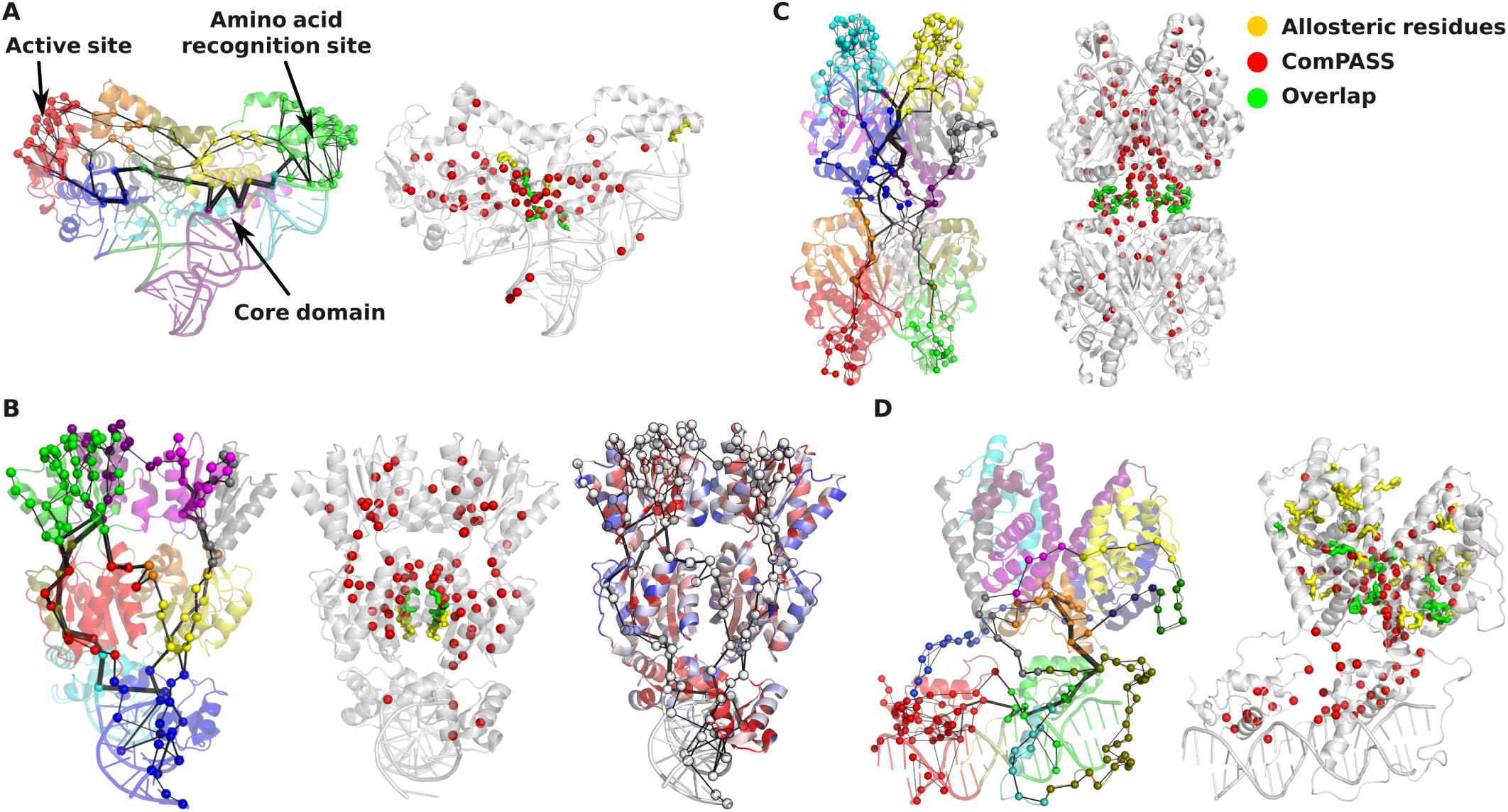
Analysis of network parameters across case study systems. The top-pathways and predicted allosteric hotspots are shown for **A)** Cysteinyl-tRNA, **B)** LacI, **C)** Bse634I and **D)** RXR*α*-LXR*β* Complexes. In all top-path figures, the thickness of the edges represents edge betweenness values, and the cartoon representations are colored based on community assignments. Spheres in red correspond to the ComPASS prediction, yellow spheres are those uniquely reported by the ASD database and green spheres correspond to the overlap between ComPASS predictions and the data from ASD. On the right panel of **B)**, the conservation profile of LacI is color coded on structure, spanning from red (highly conserved) to blue (highly variable). White spheres represent the mutational hotspots predicted by ComPASS.

We further identified the top 5% of residues based on their centrality values, which were highlighted in red (**Fig. 2**A). These residues are the most central nodes in the network, playing a crucial role in facilitating communication within the protein. Notably, a significant majority of these residues are located at the interface between the protein and RNA, underscoring their importance in mediating the communication required for efficient aminoacylation. In addition, several RNA nucleotides were also included among the predicted allosteric hotspots, further emphasizing the critical relationship between the protein and RNA in driving enzymatic activity. Among the allosteric hotspot residues, those colored in green were identified as allosteric residues in a previous study [36], while the residues marked in yellow were both identified as significant by ComPASS and previously reported as allosteric. Notably, we observed a 60% overlap between the residues identified by ComPASS and those implicated as allosteric in the prior study. This overlap highlights the consistency and reliability of ComPASS in identifying allosteric hotspots.

### Decoding Signal Transmission in LacI through Dimer Interface Pathways

In Escherichia coli, lactose metabolism is tightly regulated by the Lac repressor, a protein that blocks transcription of genes required for lactose utilization by binding to an operator region upstream of these genes. This negative regulation is alleviated in the presence of allolactose, an effector molecule that binds the repressor and triggers a conformational change, allowing gene expression [37]. The Lac repressor contains distinct functional domains: a headpiece (residues 1–49) with a helix-turn-helix motif essential for operator DNA binding, an effector binding site within the core domain (residues 62–331), and a C-terminal *α*-helix (residues 340–357) that forms a stabilizing four-helix bundle. Studying communications within the Lac repressor complex is crucial for understanding how effector molecules propagate conformational changes across its domains, enabling precise regulation of gene expression, a fundamental process in cellular function and adaptation. To elucidate these communication pathways, we applied ComPASS to MD simulations of the LacI system, providing a detailed map of the networks elucidating this regulation (**Supplementary Fig. S4** and **Fig. 2B**). The top pathways identified span across the LacI system connecting all the communities present. The DNA binding domain and the DNA together act as a community. There are two major pathways per monomer connecting the DNA binding domain (or headpiece domain) to the C-terminal helix domain. These pathways pass through the hinge domain and the core domain highlighting the role of the hinge domain in the communication network of LacI. The top-pathways surround the residues of the ligand binding region. Ligand binding might interfere with these major pathways, causing alterations in the regulation of DNA. Pathway analysis further revealed that the communication pathways connecting the ligand-binding region to DNA traverse the dimer interface, emphasizing its central role in allosteric signal transmission. Four residues have been identified as playing a pivotal role in allosteric communication of LacI [38]. Using ComPASS, we identified two of these four residues as allosteric hotspots. This result aligns with the fact that these residues were initially identified through mutational studies focused on positions critical for dimerization. The remaining predicted allosteric hotspots by ComPASS are primarily concentrated around the dimer interface. They were also found in the core domain leading up to the tetramerization domain, as well as surrounding the ligand binding region. These highlight the specific residues that play an important role in holding the dynamic network in the LacI system. Additionally, we mapped the evolutionary conservation profile of LacI from the ConSurf database on the structure, where the color code represents the conservation score (**Fig. 2B**, the right panel) [39]. From this analysis, we observed a correlation between the allosteric hotspots identified by ComPASS (white spheres on **Fig. 2B** and highly conserved residues (cartoon representations colored in red), underscoring their structural and functional importance. Additionally, comparing the top-pathways, the majority of them consist of conserved residues. These findings demonstrate the capability of ComPASS to accurately pinpoint residues critical for both the structural integrity and the regulatory function of the protein.

### Allosteric signal transmission in Bse634I

The Type IIF REase Bse634I, which recognizes the degenerate DNA sequence 5’-RCCGGY-3’ (R = purine, Y = pyrimidine), is a protein of interest because it offers a unique experimental platform for investigating the interplay between enzyme multimerization and DNA cleavage specificity. Functioning as a homotetramer arranged as a dimer of dimers, Bse634I requires the simultaneous binding of two cognate DNA copies for optimal catalytic activity [40], exemplifying an elaborate communication mechanism between its DNA-binding surfaces. The tetramer has two DNA-binding clefts, suggesting that DNA binding at one site can influence the other through allosteric effects [41]. Notably, the enzyme exhibits auto-inhibition when bound to a single DNA molecule, underscoring the regulatory cross-talk within the tetrameric assembly. This tetrameric organization stabilizes its functional dimeric unit, highlighting a sophisticated allosteric mechanism that links DNA recognition and catalytic activation. Investigating the communication and structural dynamics of Bse634I provides critical insights into the molecular basis of allosteric regulation, sequence specificity, and catalytic precision in restriction enzymes, with implications for biotechnological applications and enzymatic engineering.

After applying ComPASS to Bse634I (**Supplementary Fig. S5**), we extracted the top pathways to study the communication network in this complex (**Fig. 2C**, right panel). As the literature states and our study agrees, the interface between primary dimers plays a critical role in transmitting allosteric signals. It has been shown that mutations at this interface, such as N262A and V263A, dramatically alters the catalytic properties of Bse634I without disrupting tetramer assembly [40]. Although communication is observed across the tetramer interface, it can be regulated by the interaction between dimers. We also identified the allosteric hotspots and observed that all the residues implied in allostery (R226, W228, V262 and N263) are identified by ComPASS (**Fig. 2C**, left panel). Although, the allosteric hotspot residues are distributed across the complex, the majority of them is observed at dimer-dimer and tetramer interfaces highlighting the significance of the interfaces in communication.

### Helix5 is a Conduit for Ligand-DNA interactions in LXR*β*

We studied the retinoid X receptor *α*-liver X receptor *β* (RXR*α*-LXR*β*) heterodimer. This heterodimeric nuclear receptor complex regulates transcriptional programs crucial for lipid metabolism, cholesterol homeostasis, and inflammatory responses [42]. The heterodimer binds to specific DNA response elements to modulate gene expression, integrating cellular signaling with metabolic and immune functions. Insights into the molecular communication pathways within the RXR*α*-LXR*β* in complex with DNA, could reveal mechanisms underlying receptor activation and dysfunction, shedding light on their role in diseases such as atherosclerosis, cancer, and metabolic disorders. Using ComPASS, we extracted the communication network of RXR*α*-LXR*β* (**Supplementary Fig. S6**). **Fig. 2D** left panel shows the communication routes between the ligand binding site (colored in red) and the DNA binding site (colored in green), which are passing through the helix 5 (H5). It is known that H5 undergoes conformational changes upon ligand binding that contribute to the stabilization of the receptor’s active state [43]. Variants lacking the full ligand-binding domain, including H5, have been associated with different disease outcomes in certain cancers, highlighting the importance of this structural element [44]. In summary, and as confirmed by ComPASS, H5 is a key structural component of LXR that contributes to ligand binding, receptor activation, and cofactor recruitment. Previously, using statistical coupling analysis a set of 27 residues were proposed as energetically coupled and crucial for metabolic signaling [45]. Some of the residues among these could affect the ligand binding upon mutation to alanine. Among the mutants tested by Shulman et. al, there were namely 8 residues in RXR*α* (E239, W282, F289, D363, D379, L383, L378 and V373) that were identified by ComPASS as allosteric hotspots. The residues W282 and F289 are involved in forming the hydrophobic core of the ligand binding region, which is crucial for accommodating ligands. L383 and L378 might contribute to the hydrophobic environment of the ligand binding domain, aiding in ligand stabilization as suggested in [46]. V373 is located in the dimerization interface and may play a role in stabilizing dimer formation. D379 being involved in forming salt bridges and hydrogen bonds might contribute to dimer stability [47]. These results demonstrate the capabilities of ComPASS in identifying functionally significant residues.

### Communication in Nucleosome Complexes

The nucleosome core particle (NCP) consists of *∼*145-147 base pairs of DNA wrapped around an octamer of histone proteins, including two copies each of histones H2A, H2B, H3, and H4. In this study we were interested in understanding the differences in communication network of four different nucleosome systems namely 1KX5, 1KX5_*L*1_, 1F66, and NCP601. The 1KX5 system, serving as the control, represents a nucleosome reconstituted with an *α*-satellite DNA sequence and canonical histones. The 1KX5_*L*1_ system has mutated L1 loops and the mutations were predicted to influence the stability of the nucleosome [18]. The 1F66 system incorporates H2A.Z, a histone H2A variant, in place of canonical H2A. H2A.Z can significantly alter nucleosome dynamics and energetics. Comparing this to the canonical 1KX5 structure could provide insights into the mechanism of histone variants influence on its allosteric networks. These three systems have previously been subjected to MD simulations in a study by Bowerman and Wereszczynski [18], where they investigated the communication pathways between the L1 loop of H2A and the DNA ends and their re-shaping upon the presence of the H2A.Z variant. The fourth one, NCP601, was chosen because it harbors the 601 Widom positioning sequence instead of the *α*-satellite like in the other systems, allowing us to assess the DNA sequence effects on the communication networks within the NCP. All these systems represent different structural variations of nucleosomes, allowing for a comprehensive analysis of communication networks across diverse configurations.

In **Fig. 3**, we represent the pathways between the H2A L1 loops and the nucleotides of the DNA entry/exit site. In the 1KX5 system, the pathways originate at the L1 loop located at H2A (depicted in yellow) and pass through H2B (orange), H4 (green), H3 (blue), and finally the DNA (white). Similar pathways are observed in all other systems, although the specific dependency on subunits and their preferences varies. In Bowerman and Wereszczynski’s study, in which pathways were computed, it was demonstrated that these pathways traverse through the DNA [18]. These pathways enter the DNA at the L1-DNA interface and travel through the DNA. While the pathway through DNA could indeed be one of the routes connecting these two regions, we hypothesize that the shortest pathway might occur through the histone core itself. In the 1F66 system, in which H2A is replaced by its H2A.Z isoform, we note a decrease of the involvement of the H2B residues in the pathways linking the H2A.Z L1 loop to the DNA entry/exit site. This highlights how the communication pathways within the nucleosome are sensitive to histone variants exchange, in agreement with what was observed by Bowerman and Wereszczynski [18]. In the case of NCP601, we observed a slight increase in DNA involvement within the communication pathways, suggesting that the 601 Widom sequence may subtly influence nucleosome stability. This finding indicate a potential contribution of sequence-specific dynamics to nucleosome organization. Interestingly, H3Y54 and H3S57, that are well-known phosphorylation sites involved in gene transcription and DNA repair processes [48, 49], are consistently present in the shortest pathway from H2A L1 loop to the DNA entry/exit point, further supporting their crucial role in the nucleosome architecture modulation.

**Figure 3:**
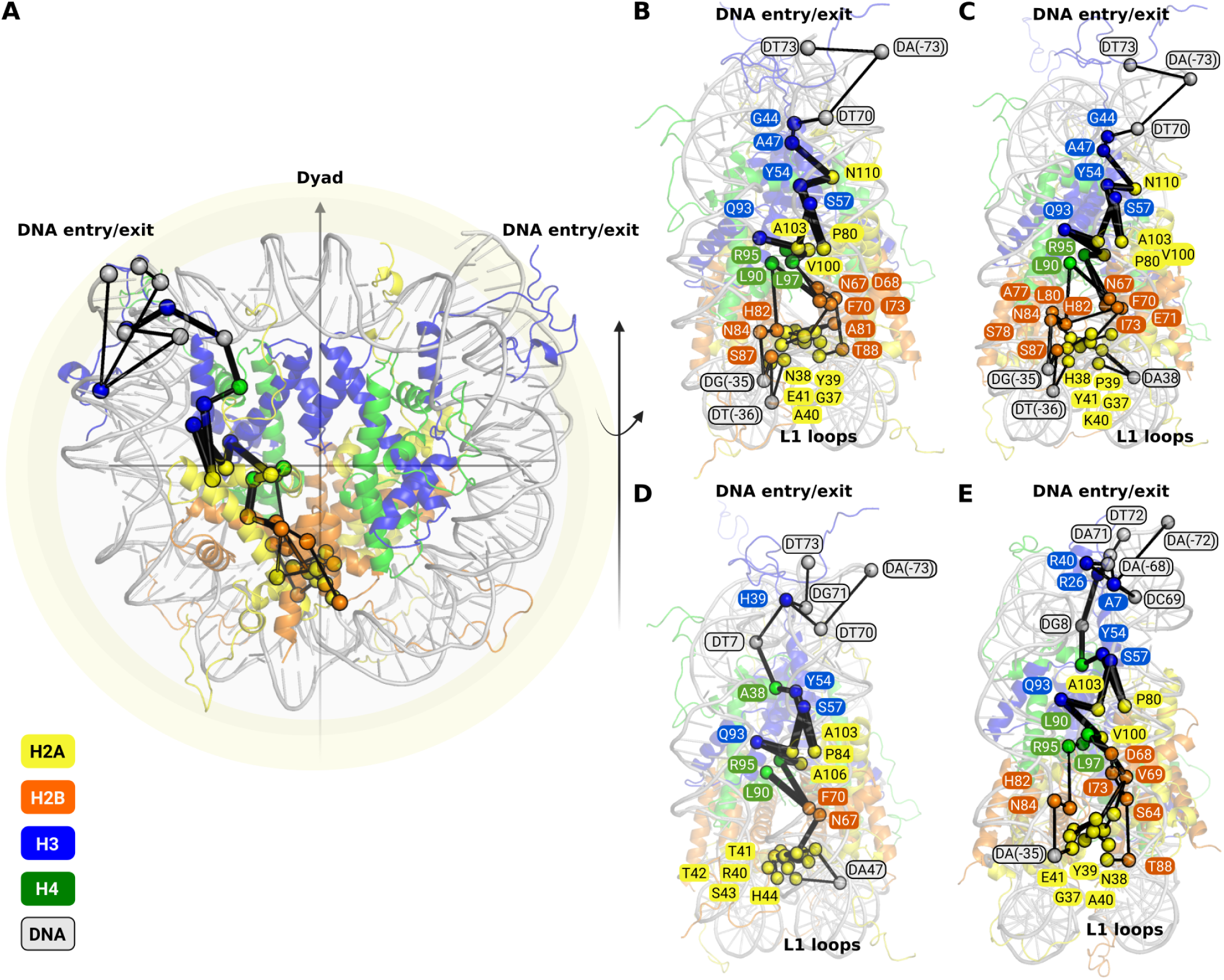
Comparison of L1-DNA entry/exit site pathways across the four nucleosome systems. **A)** A general representation of the pathways identified between the L1 loop and the DNA entry/exit site. Specific pathways are highlighted for each nucleosome system: **B)** 1KX5, **C)** 1KX5_*L*1_, **D)** 1F66, and **E)** NCP601. In each complex, the histone proteins are color-coded as follows: H2A (yellow), H2B (orange), H4 (green), and H3 (blue). Computed pathways are visualized as lines connecting residues, which are represented as spheres. Panels on the right show the nucleosome complexes after a 90-degree rotation, with labeled residues providing a clearer depiction of the pathways in each system.

We conducted a community analysis revealing that all nucleosomes form dynamic communities that deviate significantly from structural domains. These dynamic communities are predominantly governed by the spatial orientation of residues (**Supplementary Fig. S7**). Notably, DNA regions in contact with histone proteins consistently form integrated communities with the protein components. In the 1KX5_*L*1_ nucleosome, the *α*1 helix of histone H2A, along with the DNA at the entry/exit site, forms an independent community. This observation suggests that the mutation of the L1 loop may influence community formation by enhancing the role of the H2A *α*1 helix. In contrast, in the NCP601 system, the DNA entry/exit site forms a separate community on its own. Furthermore, we observed that in NCP601, only a single round of DNA integrates into the protein community, likely due to the distinct DNA turn in this system compared to the other nucleosome complexes. We compared identified communities with the communication blocks detected by COMMA reported previously [50]. Although COMMA identified fifteen communication blocks within the histone core, ComPASS detected ten communities in the histone core of 1KX5 (**Supplementary Fig. S8**). Interestingly, the communities identified by ComPASS were more regionally localized and encompassed all proteins within specific structural regions, whereas the communication blocks detected by COMMA were largely confined to H2A-H2B and H3-H4 dimers, with limited extension into the broader structural framework. This divergence likely arises from methodological differences, including the incorporation of nucleotides in the ComPASS analysis and the application of a distance cutoff to define the residue-residue network. This approach in ComPASS appears to result in more regionally specific communities, in contrast to the broader communication blocks observed with COMMA. Despite these differences, both methodologies revealed consistent per-dimer distributions, with each community or communication block predominantly composed of H2A-H2B or H3-H4 pairs. Histones H3 appear to play a major role in the communication network spanning the upper part of the nucleosome core particle, which is in line with the COMMA analysis of the histone core we described in a previous work [50]. Importantly, the identified top paths bridging nucleic acids and protein residues are highly present at the dyad (**Supplementary Fig. S7**), which agrees with the fact that the histone-DNA interaction are especially strong in this region.

### Comparison with other existing methods

We compared the results of ComPASS with other available methods, including NRIMD, MDiGest, and PyInteraph2.0 for the prediction of allosteric pathways in the case of CysRS complex. Previously, using MD simulations a pathway has been established between residues in the anticodon-binding domain and the active site of CysRS [36]. This pathway comprises five intervening residues, deduced based on high correlation values between the anticodon-binding domain and the active site. Our findings reveal overlapping and unique features across the methods as shown in **Fig. 4**. While some residues are common to the pathways identified by multiple approaches, ComPASS stands out as the only tool to include nucleotides in the pathway. NRIMD predicted pathways involving residues unexpectedly separated by distances ranging from 15 to 30 Å. The NRIMD webserver is also limited by the trajectory length and size constraints requiring individual residue pairs to be specified for pathway calculations, which complicate multiple pathways analysis. On the other hand, MDiGest is restricted by its reliance on a single property among either distances or dihedrals, or Kullback–Leibler divergence. PyInteraph2.0 predicts pathways with some residues overlapping those reported in [36]. Among all methods, ComPASS emerges as a comprehensive approach, integrating multiple metrics to provide feasible and biologically relevant pathways. By leveraging distance criteria, ComPASS computes pathways that align closely with expected structural and functional dynamics, offering a plausible mechanism of communication between the anticodon-binding domain and the active site. While we acknowledge that the computed pathway is one of several possible routes, ComPASS provides a robust framework for identifying and analyzing communication pathways in protein-NA complexes.

**Figure 4:**
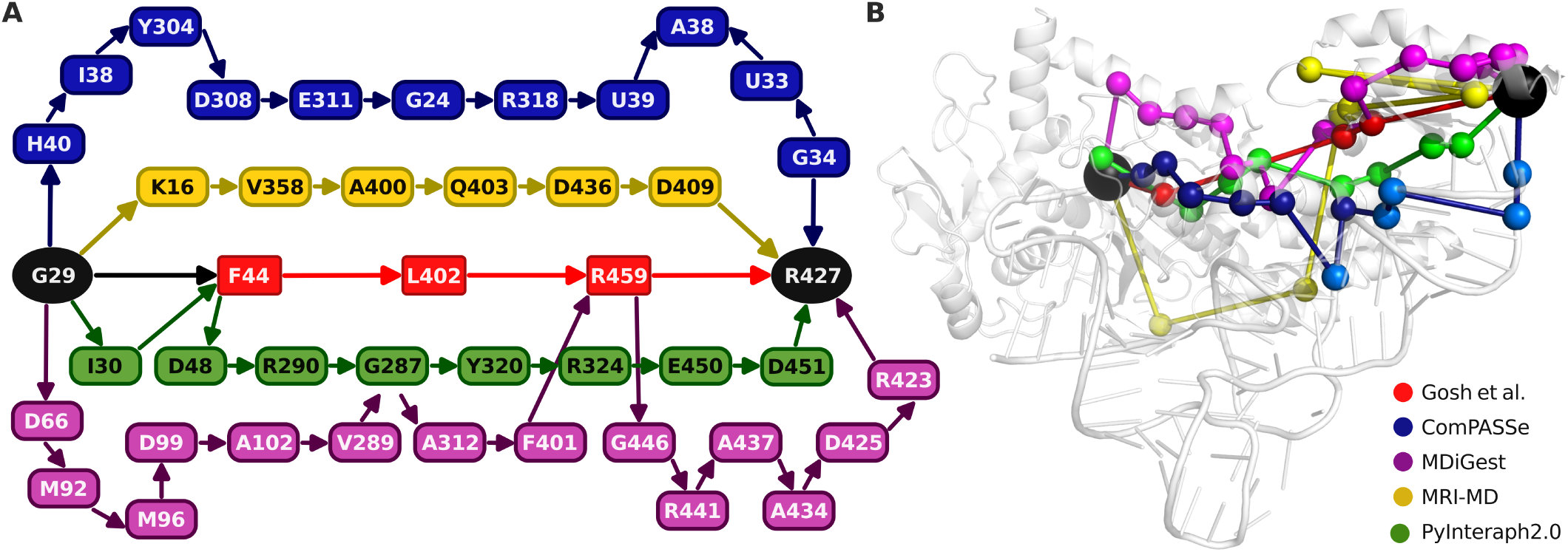
Comparison of communication pathways of CysRS complex computed using Com-PASS, MDigest, PyInteraph2.0 and NRI-MD. Pathways were computed between residues G29 and residue R427. On the left, the pathways are represented as a network diagram, where the source (G29) and target (R427) are colored in black. On the right, the pathways are mapped onto the structure of the CysRS. The pathway reported in Gosh et al. [36] is shown in red, and the pathways computed by NRI-MD, MDiGest, PyInteraph2.0 and ComPASS are represented in yellow, purple, green and blue, respectively. In the case of ComPASS we used dark and light shades of blue to distinguish amino acids from nucleotides, respectively.

## Discussion

Communication networks are fundamental to the function of protein-NA complexes, driving long-range signaling and regulation of critical biological processes. In these systems, allosteric signaling is intricately linked to structural remodeling, enabling the transmission of signals from one site to a distant functional site. For instance, in the CRISPR-Cas9 complex, binding to the PAM recognition sequence induces coupled motions within the protein framework, triggering conformational shifts that activate the system for precise and concerted DNA cleavage [51]. Such mechanisms underscore the importance of understanding communication and allostery in protein-NA complexes, as they play pivotal roles in regulating processes like DNA/RNA replication, chromatin remodeling, and gene editing—offering immense potential for applications in medicine and bioengineering. Despite the recognized importance of allostery in these complexes, there is a scarcity of computational tools specifically designed to study these phenomena in protein-NA systems. This gap is addressed by ComPASS, a novel method that provides a comprehensive framework for analyzing communication pathways in macromolecular complexes using MD data. By integrating diverse properties such as communication propensity, generalized correlations, distances, and interaction strengths, ComPASS offers a holistic approach to mapping communication networks, revealing dynamic pathways that underlie the functional mechanisms within both protein-protein and protein-NA complexes.

In comparison to existing methods, ComPASS demonstrates a significant performance in identifying realistic communication pathways within macromolecular complexes. Notably, pathways proposed by some methods often include edges spanning distances of *≃*30 Å, which may not represent practical or biologically plausible pathways. In contrast, ComPASS identifies residues in close proximity, delineating feasible communication routes supported by structural and dynamic considerations. This advantage can be attributed to ComPASS’s ability to integrate multiple properties rather than relying on a single metric. A broader challenge in this field remains the scarcity of experimentally validated data on allostery. Experimentally confirming allosteric residues is inherently difficult, often limiting validation to a subset of residues predicted by computational methods. Despite this, ComPASS achieves high accuracy, identifying known allosteric residues reported in the literature with success rates ranging from 30% to 100%, depending on the system. This variability highlights the complementary roles of computational and experimental approaches, as experimental techniques can only confirm the role of a few residues within the predicted network. Beyond specific cases, ComPASS also enables the establishment of communication networks in complex systems such as nucleosomes, allowing for comparative analyses of different nucleosome complexes. These findings underscore the potential of ComPASS to uncover nuanced differences in communication dynamics, providing a powerful tool to study the intricate mechanisms of macromolecular systems.

ComPASS is a versatile and efficient tool for studying macromolecular communication networks. First, as a Python package, it offers significant advantages over web-based platforms, which often impose limitations on system size and computational resources, particularly for handling long and resource-intensive MD trajectories. Second, its open-source architecture allows users to customize and adapt the code, facilitating the modification of various properties to meet specific research requirements. Third, the integration of Numba just-in-time compilation ensures exceptional computational performance, enabling ComPASS to surpass existing methods in both speed and efficiency. **Supplementary Table S2** reports the performance of ComPASS on all the case studies. The computations were performed on a machine equipped with an 126 GB of RAM and an Intel Xeon Silver 4216 CPU (2.10 GHz) using 32 threads. Moreover, ComPASS provides a comprehensive platform for analyzing communication within macromolecular complexes, particularly those involving nucleic acids. It enables users to identify critical residue interactions, compute shortest communication pathways, and visualize intricate communication networks in an intuitive manner. Finally, ComPASS supports the analysis of multiple MD replicas, offering a robust framework to capture the dynamic behavior of biomolecular systems. These capabilities position ComPASS as a powerful tool for unraveling complex biological phenomena, advancing our understanding of macromolecular dynamics, and aiding in the rational design of targeted therapeutics.

## Data availability

The ComPASS code is freely available at https://github.com/yasamankarami/compass. All the MD trajectories and final output of the case studied have been deposited in the Zenodo database (accession doi:10.5281/zenodo.14843518).

## Acknowledgments

This work was granted access to the HPC resources of IDRIS under the allocation 2023-A0150714660 (granted to Y.K.) and 2023-A0150714577 (granted to E.B.) made by GENCI.

