## Supplementary Information for "Communication pathway analysis within protein-nucleic acid complexes"

### Supplementary Methods

This section provides additional details on the methodology used in the ComPASS package.

#### Execution of ComPASS and input parameters

To run the ComPASS method, the user must provide a configuration file that includes essential details such as trajectory files, distance cutoff values for graph construction, output directory, and an option to calculate optimal communication pathways. The configuration file serves as the primary interface for user input, allowing customization of the parameters that govern the network construction and analysis processes. The trajectory files, typically containing molecular dynamics data, are required to generate the corresponding communication networks, with specific cutoffs that dictate the edge formation criteria based on distances between residues. The output folder specified by the user will store the results of all computations.

#### Output files and interpretation of results

Upon execution, ComPASS generates a primary output file and two subfolders: matrices and network. The primary output file contains detailed mappings between the user-provided PDB file and the corresponding graph construction process, ensuring traceability of the data used to build the communication network.

The matrices subfolder stores all the matrices discussed in the methods section, each provided in two formats: text files for the matrices (.mat) and the corresponding heat map images (.png) for visualization. These matrices include the generalized correlation matrix, interaction frequency matrix, and communication propensity matrix, which are essential to understand residue interactions and network construction.

The network subfolder contains several files designed to assist with the visualization and interpretation of the constructed communication network. These files include various graph representations in the form of PyMOL files (.pml), enabling 3D visualization of the network structure. Additionally, the folder includes several text files that describe the graph properties, such as centrality measures, community assignments, and edge betweenness data. For instance, the community assignments are provided in both raw text and visual formats, indicating which residues belong to each functional module. Edge betweenness data highlights the importance of specific connections in facilitating communication between residues within the network.

Furthermore, the folder includes the results of shortest path and alternative path computations, allowing users to examine communication pathways between residues. The file containing shortest path data provides a detailed list of the top paths between residue pairs, ranked by their communication efficiency. Histogram plots and JSON files offer additional data visualizations and graph data representations, while PML files for different cutoff values provide alternative views of the graph at varying thresholds, giving users flexibility in their analysis. All of these output files work together to offer a comprehensive overview of the communication network, aiding in the interpretation of protein-RNA interactions and the identification of key regulatory residues.

### Supplementary results

#### Cysteinyl-tRNA

The network parameters cliques and communities are valuable tools for capturing subtle conformational changes in protein structures, particularly in the context of ligand interactions, as they are often associated with regions of local rigidity or flexibility. In the case of the CysRS system, we applied network analysis to identify a total of 29 cliques across the entire structure. These cliques, which are densely connected subgraphs where each node is connected to every other node, are key to understanding how different regions of the protein communicate and adapt during the tRNA recognition process. Among these cliques, three were particularly notable for their functional significance, as illustrated in **Supplementary Fig. S3B**. Clique 1, located in the domain critical for anticodon recognition, is essential for the enzyme's ability to properly attach the amino acid to the tRNA by recognizing the anticodon. This region is crucial for ensuring the specificity of the aminoacylation process. Clique 2 corresponds to the CP loop, which is known to undergo conformational changes upon tRNA binding. These changes are necessary for the proper functioning of the enzyme and the efficiency of the aminoacylation reaction. Clique 3 is associated with the helical arm of the complex, which, in conjunction with the A10.U25.U45 region, forms a vertical arm structure. This structural feature compensates for the lack of tight contacts at the inner elbow of the protein, thus stabilizing the protein-RNA interface. These three cliques highlight key regions of the protein involved in tRNA recognition and interaction, demonstrating the dynamic nature of the CysRS-tRNA complex. In addition to cliques, we also identified several communities within the network. These communities differ significantly from the structural and functional domains of the protein. For instance, the Rossmann fold and CP domain, together with RNA, form a distinct community that ensures the proper embedding of the RNA tail into the protein structure. As shown in **Supplementary Fig. S3C**, many of the communities span both protein and RNA components, reflecting the intricate interplay between the two entities during the tRNA recognition process.

#### LacI

The signal transduction in LacI occurs between two functional regions: the inducer-binding site and the DNA-binding region, with contributions from the dimerization interface. In **Supplementary Fig. S4A**, we show that residues in these regions form distinct and independent cliques. Furthermore, **Supplementary Fig. S4B** highlights the predicted community structure within LacI, revealing functionally significant groupings of residues. Notably, these communities suggest compensatory effects, where mutations in one residue might be buffered by residues within the same community, although this hypothesis requires experimental validation. Finally, in **Supplementary Fig. S4C**, we present the putative communication pathways within LacI, which confirm asymmetric signaling and cross-monomer interactions. These routes propagate the signal from the inducer-binding site of one monomer to the DNA-binding site of the other through a network of interconnected residues, consistent with prior predictions [1]. Together, these findings offer a detailed view of LacI's communication dynamics, providing a framework for future experimental studies and applications in synthetic biology.

#### Bse634I

In the restriction enzyme Bse634I, two distinct communication signals govern its activity: an inhibitory "stopper" signal and an activating "sync" signal. These signals propagate through the dimer-dimer interface, with their interplay shaping the enzyme's catalytic and regulatory properties. As shown in **Supplementary Fig. S5A**, three cliques were identified across the entire complex, with residues at the dimer-dimer interface forming a distinct clique (Clique 2). Additional cliques were observed at the DNA-binding site and the dimer interface, underscoring the functional significance of these regions. Community analysis (**Supplementary Fig. S5B**) further revealed three communities per subunit, with the third community consistently located at the dimer-dimer interface, encompassing loops that mediate critical inter-dimer interactions. Mapping the communication network (**Supplementary Fig. S5C**) revealed a striking asymmetry, where communication signals are predominantly biased toward one dimer. This observation aligns with previous studies suggesting that the allosteric signal originates asymmetrically in one monomer and propagates to the other through non-covalent interactions [2]. Together, these findings indicate a detailed communication map of Bse634I, highlighting the central role of the dimer-dimer interface in mediating both inhibitory and activating signals. The observed asymmetry in signal propagation offers a mechanistic explanation for the enzyme's allosteric regulation, suggesting that inter-dimer communication might modulate its catalytic efficiency and regulatory precision. These insights pave the way for further investigations into the functional relevance of asymmetric signaling in similar systems.

#### RXR$\alpha$-LXR$\beta$ complex

Ligand binding to one partner in the RXR$\alpha$-LXR$\beta$ complex induces conformational changes that propagate through the heterodimer interface, modulating the activity of the other partner. This mechanism allows the complex to integrate multiple ligand signals and finely regulate transcriptional outcomes. In **Supplementary Fig. S6A**, we identified distinct cliques within the complex, with a notable clique encompassing residues at the DNA interface, indicating their central role in signal transduction. Community analysis (**Supplementary Fig. S6B**) revealed four primary communities per subunit, with the third community consistently localized at the heterodimer interface, comprising residues critical for cross-subunit communication and another belonging to DNA binding. The comprehensive communication network, visualized in **Supplementary Fig. S6C**, highlights the role of hinge domain in signal propagation, with communication routes prominent along the hinge domain connecting DNA binding domain to the regulatory domain. This finding aligns with previous studies, which suggest that RXR$\alpha$ serves as the primary integrator of ligand signals[3], with its conformational shifts directing the activity of LXR$\beta$ through non-covalent interactions. Together, these results provide a detailed communication map of the RXR$\alpha$-LXR$\beta$ complex, underscoring the pivotal role of the heterodimer interface in facilitating coordinated activity. The observed asymmetry suggests a regulatory hierarchy within the complex, where RXR$\alpha$-driven signals fine-tune LXR$\beta$ activity, enabling dynamic transcriptional control in response to diverse signaling cues.

#### Nucleosome studies

Allosteric hotspots, defined as key residues critical for mediating long-range communication and regulation, exhibit system-specific distributions that underscore distinct regulatory mechanisms within each system as shown in **Supplementary Fig. S7**. In 1KX5$_{L1}$, the allosteric hotspots were predominantly dispersed along the DNA-protein interface, distinguishing this system from others. Notably, the H2A L1 loops consistently harbor allosteric hotspots across all four systems, emphasizing their functional significance in facilitating allosteric communication. In 1F66, hotspots cluster primarily around the H3-H2A.Z interface, underscoring the regulatory role of the H2A.Z variant. Across systems, hotspots are most prominent on the DNA entry/exit sites and at the dyad (at the center of the DNA sequence), suggesting a conserved mechanism for DNA engagement. Comparative analysis of the network architectures reveals that protein-DNA interactions are integral to network organization, as evident from the pronounced connectivity between protein and DNA regions.

### Supplementary Tables

Table S1: **The list of performed MD simulations.** All studied systems, as well as number of replicates and simulation time are reported here.

| System | Simulations length (ns) | #Replicates | #Atoms | Box size (Å$^3$) |
|---|---|---|---|---|
| CysRS | 500 | 3 | 136,838 | 1,340,689 |
| LacI | 500 | 3 | 121,304 | 1,154,828 |
| Bse634I | 500 | 3 | 248,754 | 2,400,957 |
| RXR$\alpha$-LXR$\beta$ | 500 | 3 | 164,846 | 1,592,945 |
| 1KX5 | 250 | 3 | 231,078 | 2,559,465 (oct) |
| 1KX5$_{L1}$ | 250 | 3 | 234,708 | 2,596,785 (oct) |
| 1F66 | 250 | 3 | 234,624 | 2,598,607 (oct) |
| NCP601 | 5000 | 3 | 365,854 | 3,993,490 (oct) |

Table S2: **Overview of ComPASS run time.** The table reports wall time required to execute ComPASS on each case study using 32 CPU cores.

| System | #Residues | #Atoms | Number of frames | Wall Clock Time (sec) | System Time (sec) |
|---|---|---|---|---|---|
| CysRS | 533 | 9515 | 15,003 | 169.68 | 69.81 |
| LacI | 692 | 11085 | 15,003 | 272.38 | 130.52 |
| Bse634I | 1204 | 20485 | 15,003 | 639.68 | 421.78 |
| RXR$\alpha$-LXR$\beta$ | 776 | 12890 | 15,003 | 315.55 | 130.83 |
| 1KX5 | 1268 | 25090 | 603 | 147.59 | 75.44 |
| 1KX5$_{L1}$ | 1268 | 25120 | 603 | 161.75 | 75.82 |
| 1F66 | 1266 | 24930 | 603 | 170.68 | 61.01 |
| NCP601 | 1262 | 26135 | 4525 | 311.77 | 196.57 |

Table S3: **Comparison of performance and remarks on methods used to evaluate results.** The table reports execution time and remarks for running the mentioned methods, except for NRI-MD (web-server), on the CysRS system using a 32-core system.

| **Method** | **Time (seconds)** |
|---|---|
| ComPASS | 169.68 |
| MDiGest | 670.65 |
| NRI-MD | ∼500 |
| PyInteraph2.0 | ∼820 |

### Supplementary Figures

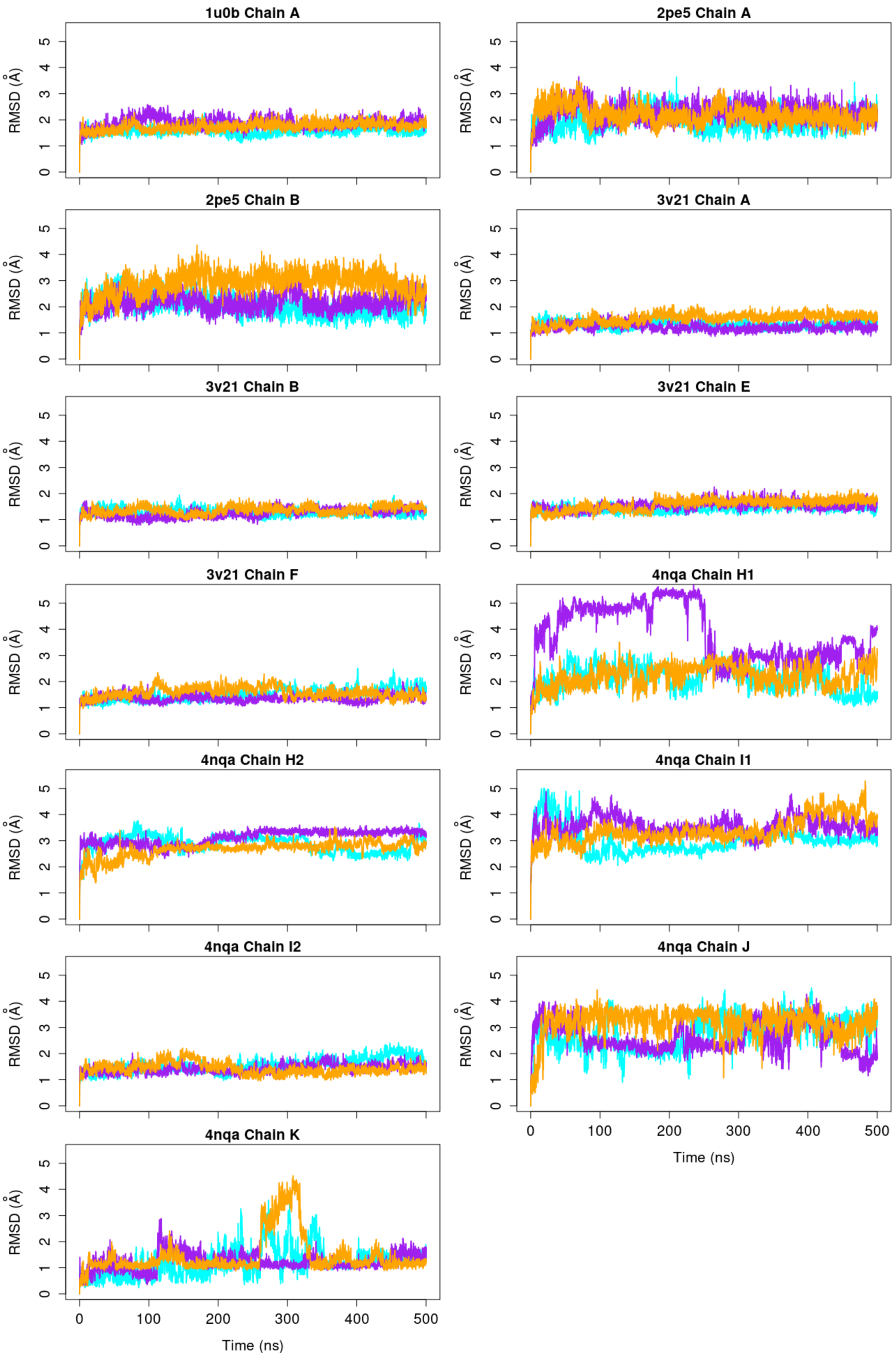


Figure S1: **Root mean square deviations of every chain for each system over MD simulations.** The RMSD values are reported with respect to the equilibrated conformation and measured over C$\alpha$ atoms for every chain of CysRS (PDB codes: 1u0b), LacI (PDB code: 2pe5), Bse634I (PDB code: 3v32) and RXR$\alpha$-LXR$\beta$ (PDB code: 4nqa). The different colors of cyan, purple and orange represent results for one of the three replicates. In case of RXR$\alpha$-LXR$\beta$, the RMSD of chains H and I are measured over the two individual domains that are connected by a linker.

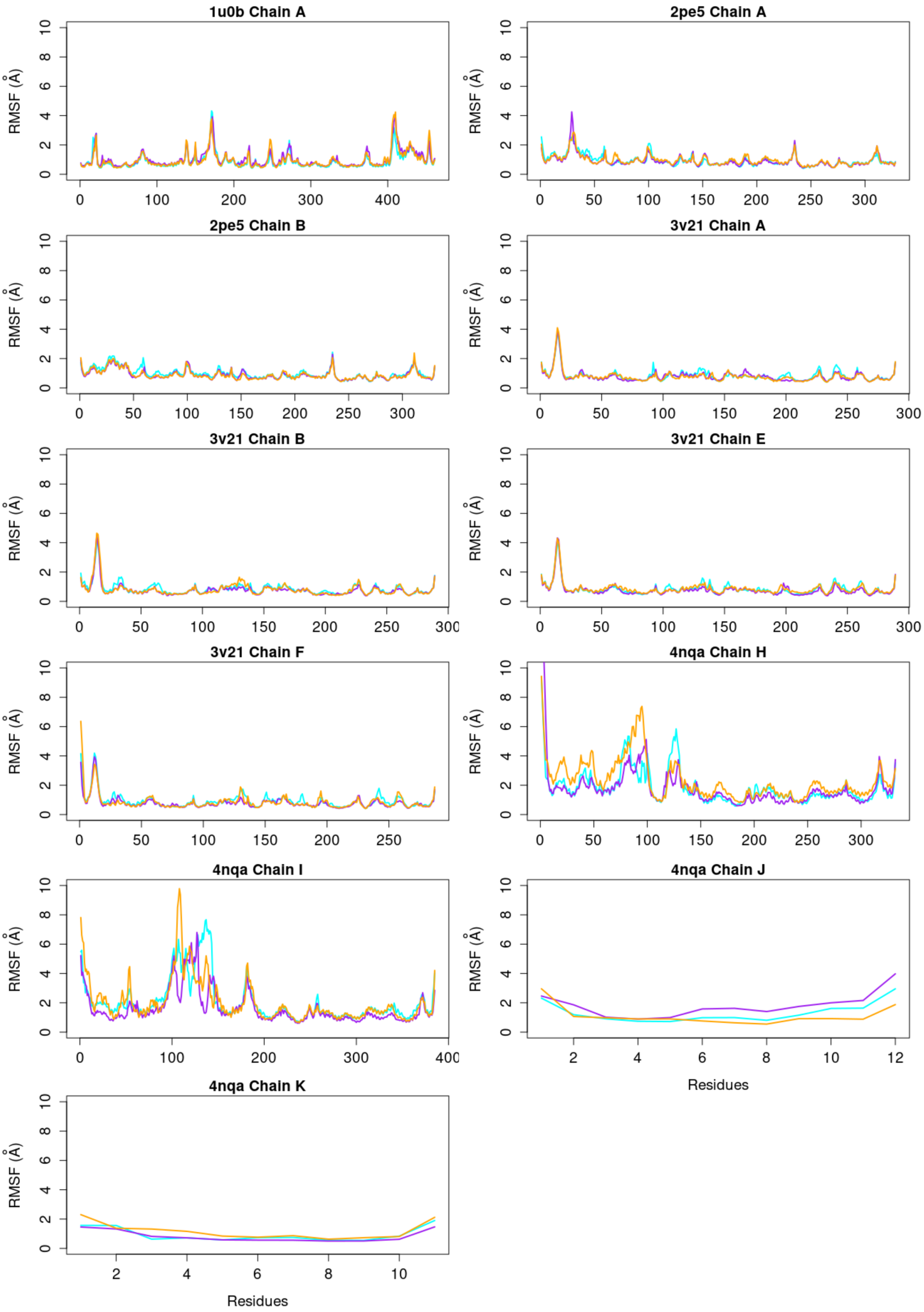


Figure S2: **Root mean square fluctuations of every chain for each system over MD simulations.** The RMSF values are measured with respect to the equilibrated conformation and over C$\alpha$ atoms of each residue. The results are reported for every chain of CysRS (PDB codes: 1u0b), LacI (PDB code: 2pe5), Bse634I (PDB code: 3v32) and RXR$\alpha$-LXR$\beta$ (PDB code: 4nqa). The different colors of cyan, purple and orange represent results for one of the three replicates.

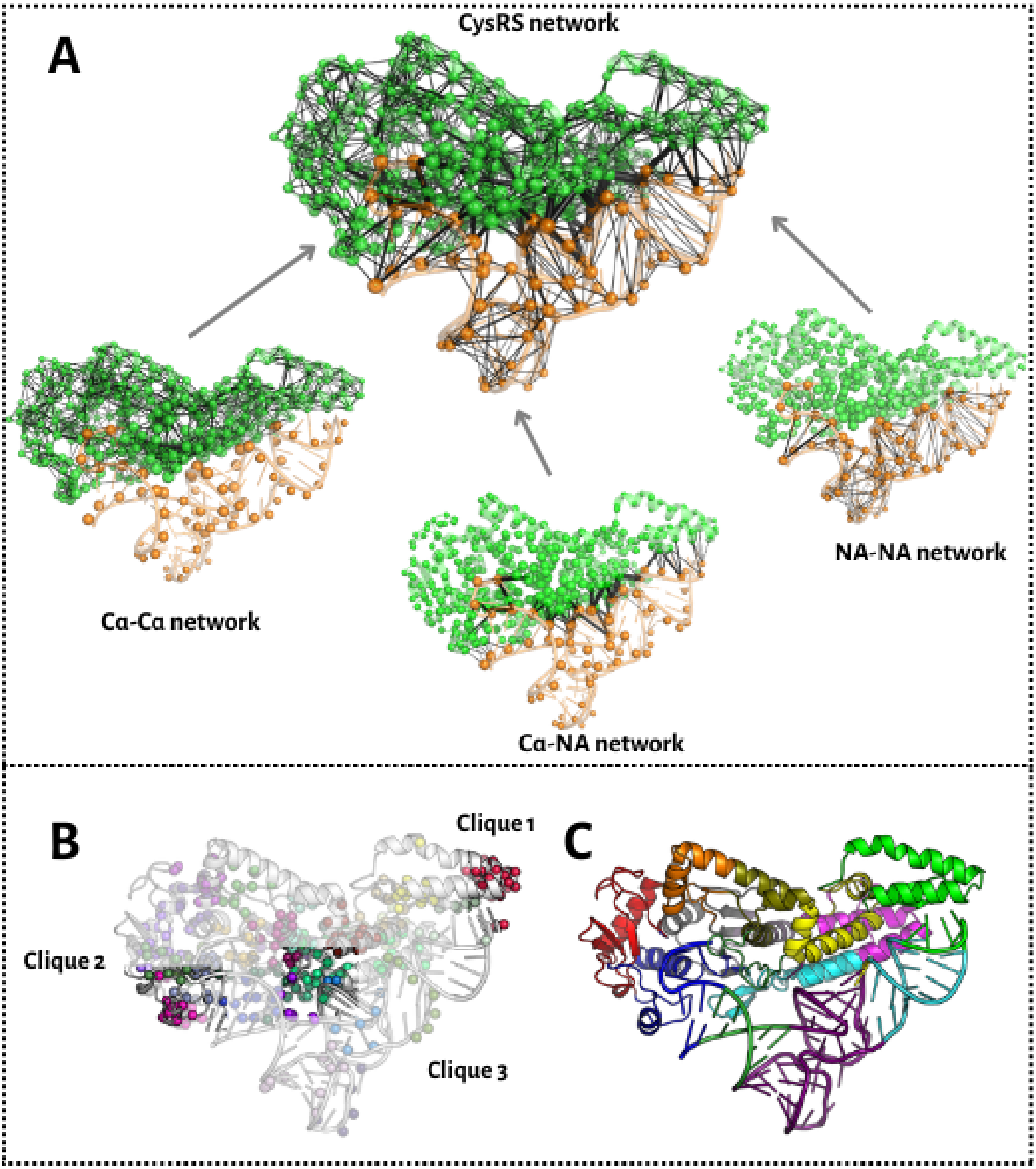


Figure S3: **Analysis of various network parameters of Cysteinyl-tRNA complex.** (A) The entire network constructed from the cysteinyl-tRNA system. As shown, the network comprises Cα-Cα, Cα-NA, and NA-NA networks, where NA refers to nucleic acid. (B) Computed cliques and communities in the cysteinyl-tRNA complex.

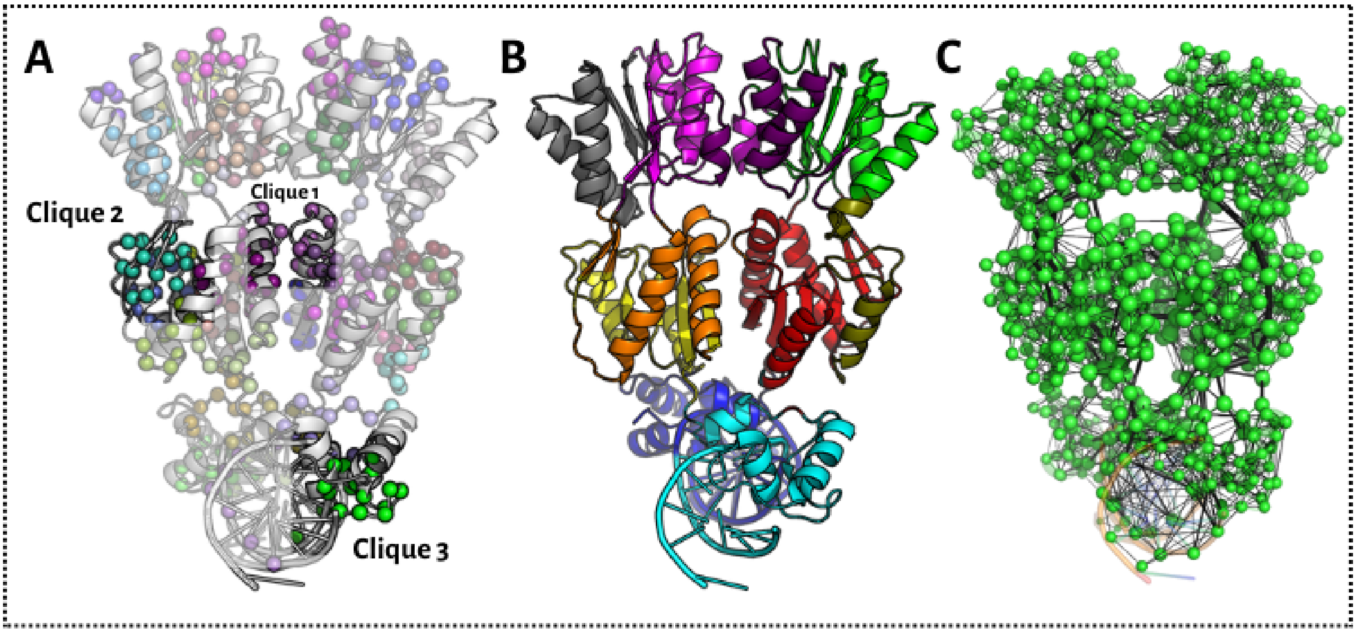


Figure S4: **Analysis of various network parameters of LacI complex.** (A) Computed cliques (B) Communities and (C) Entire network.

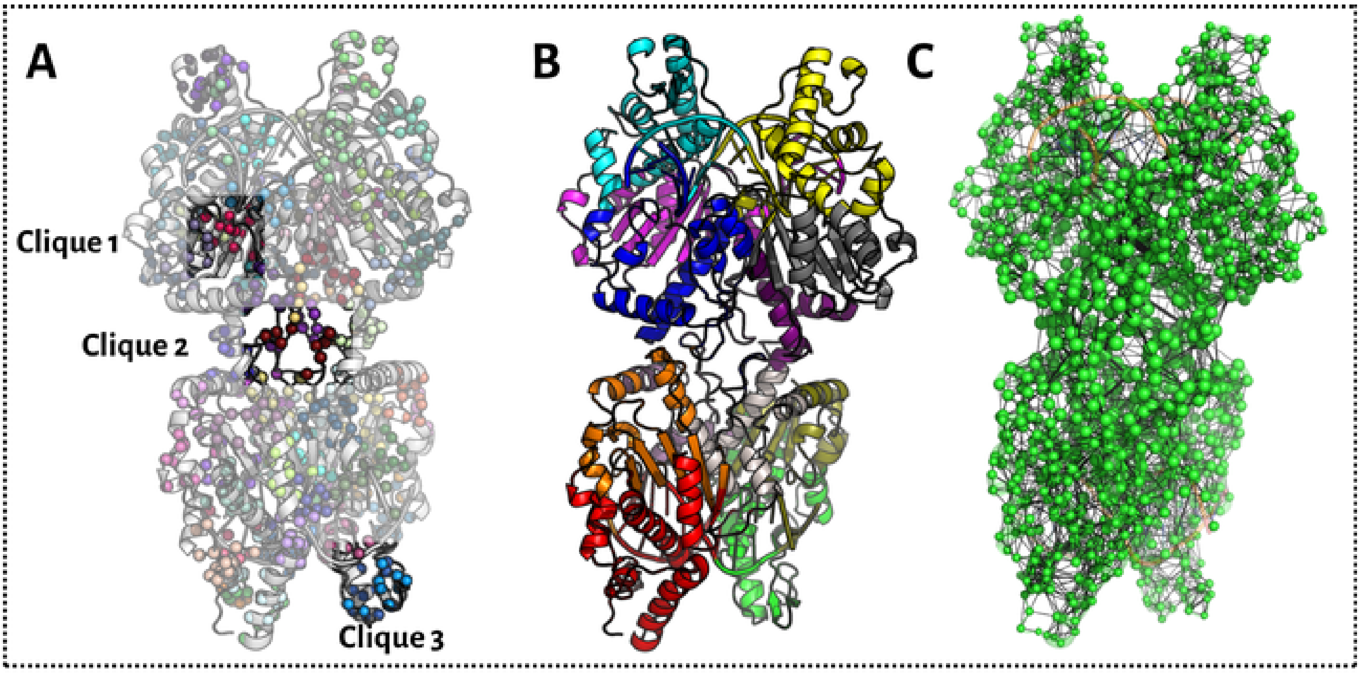


Figure S5: **Analysis of various network parameters of Bse634I complex.** (A) Computed cliques (B) Communities and (C) Entire network.

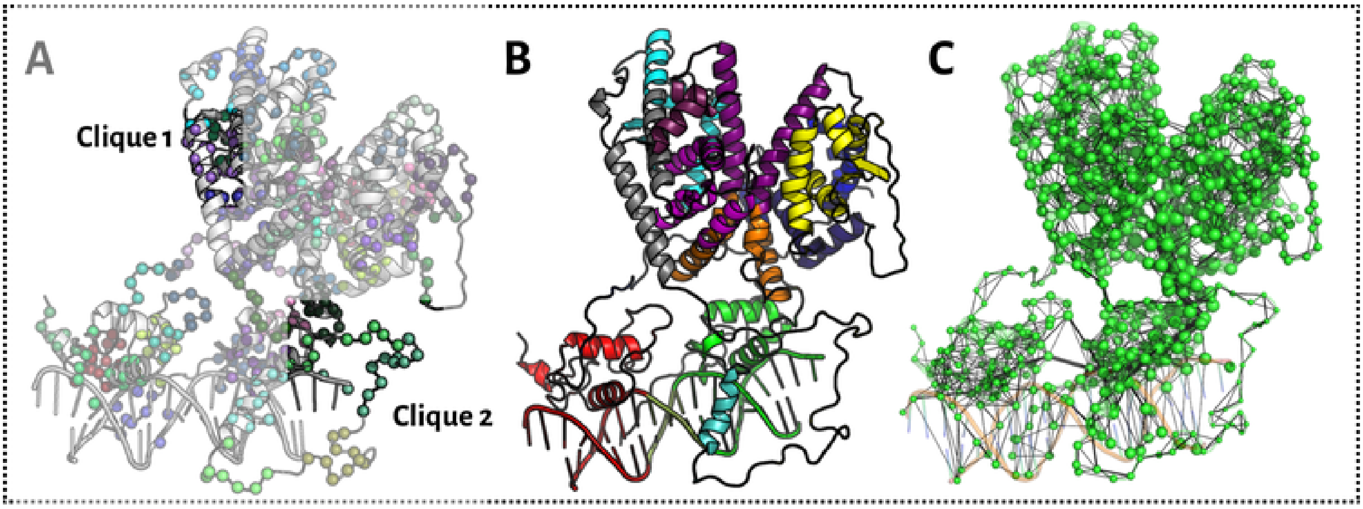


Figure S6: **Analysis of various network parameters of RXR$\alpha$-LXR$\beta$ complex.** (A) Computed cliques (B) Communities and (C) Entire network.

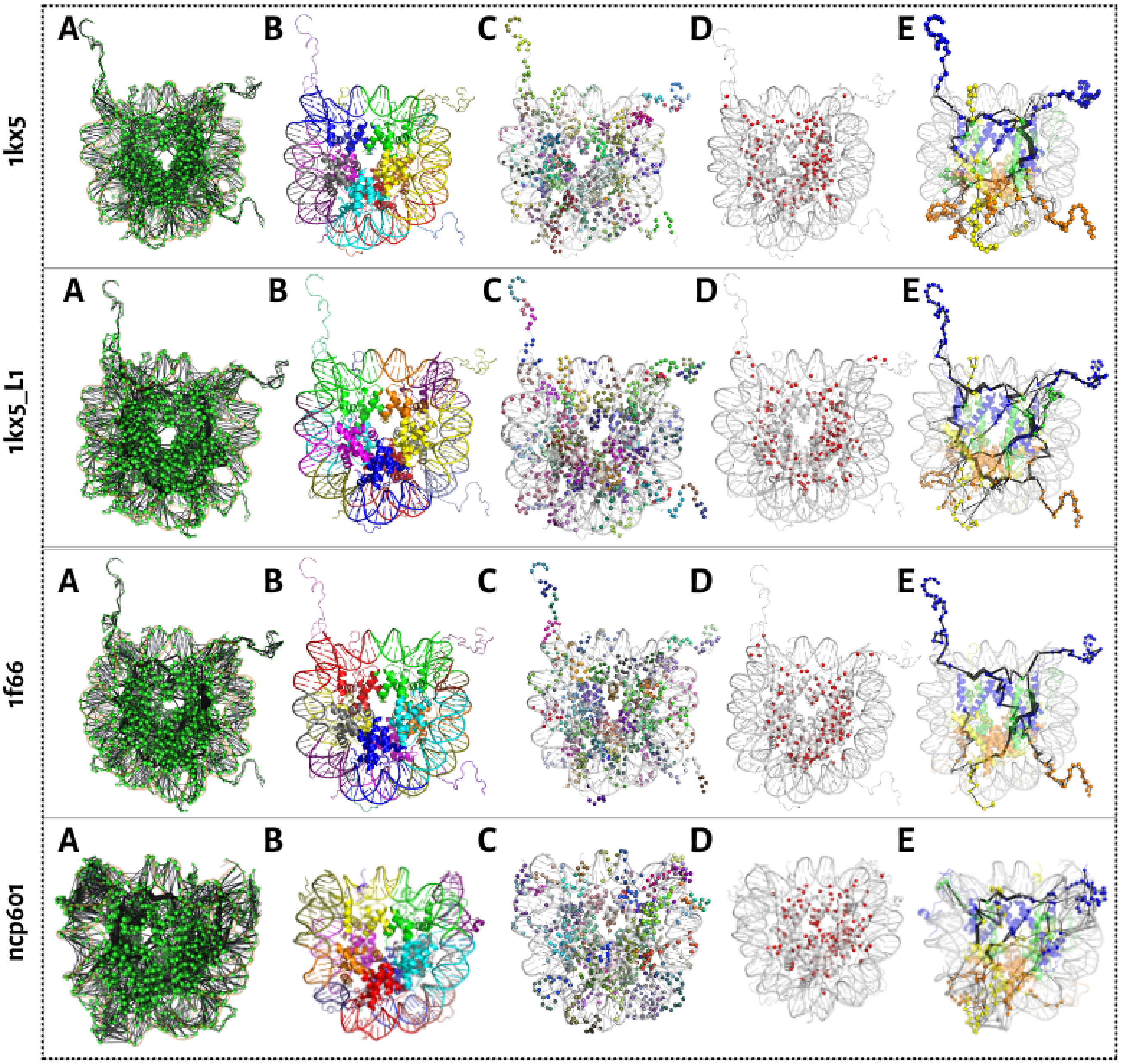


Figure S7: **Analysis of various network parameters of nucleosome complexes.** The names of the complexes are labelled in the figure. (A) Entire network (B) Communities and (C) Computed cliques (D) Hotspots and (E) Top paths where the cartoon is colored according to the nature of the protein.

molecule

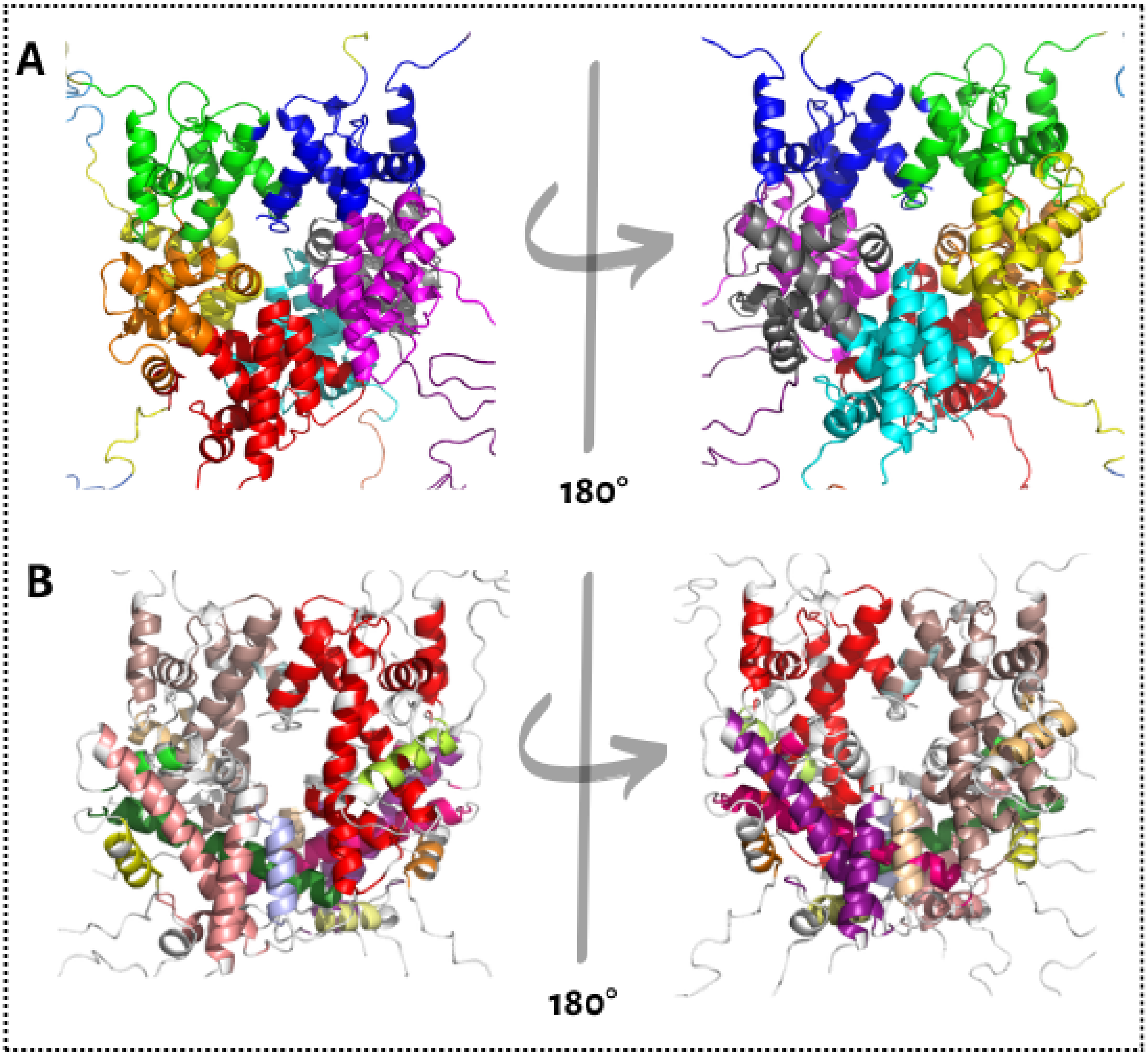


Figure S8: **Comparison of communication blocks computed from COMMA with communities in ComPASS A.** Communities calculated using ComPASS. On the right, we turned it 180 degrees. **B.** Communicaion blocks obtained using COMMA [4].
